# Wetland birds in the northern prairie pothole region may show sensitivity to agriculture

**DOI:** 10.1101/2020.09.15.298265

**Authors:** Jody Daniel, Heather Polan, Rebecca C Rooney

## Abstract

Wetland losses in the Northern Prairie Pothole Region (NPPR) are largely attributed to agriculture. Since land-use is known to influence bird habitat selection, bird community composition is likely sensitive to the extent of neighboring agricultural activity. We determined which local and landscape habitat variables are most predictive of wetland bird assemblage occurrence in southern Alberta. We:1) identified distinct bird assemblages with a cluster analysis, 2) identified which species were indicative of these assemblages using an indicator species analysis and 3) predicted which bird assemblage would occur in a wetland with a classification and regression tree. Avian assemblages were more loosely defined and had few indicator species. Importantly, assemblages were specific to the natural region in which the wetland occurred. Also, landscapes with higher agricultural activity generally supported waterfowl and shorebirds, likely because agricultural activities excluded wetland-dependent birds that nest in upland habitat. Though waterfowl and shorebirds show poor sensitivity to surrounding landscape composition, edge-nesting wetland avifauna may make good indicators of ecological integrity.

## Introduction

The majority of wetland losses in the Northern Prairie and Parkland Region (NPPR) are attributed to agricultural and urban development (Kennedy and Mayer 2002; Mitsch and Gosselink 2015), with agriculture leading to losses of about 90% of historic wetlands by 1951 in the Canadian NPPR (Bethke and Nudds 1995). Wetlands lost to agriculture are usually filled or drained to protect neighboring croplands from flooding and to increase cropland area (Schindler and Donahue 2006; Verhoeven and Setter 2010). The remaining wetlands undergo physical and chemical alterations (Rashford et al. 2011), which include 1) increased sedimentation due to tillage (Zedler and Kercher 2005) and livestock grazing (Bloom et al. 2013); 2) higher nutrient loading from fertilizer use (Schindler and Donahue 2006); 3) slower recovery rates when exposed to disturbance (Bartzen et al. 2010); and 4) lengthened hydropderiods as soil infiltration rates are lowered (van Der Kamp et al. 1999) and runoff is consolidated (McCauley et al. 2015). Thus, wetlands that have escaped drainage or infilling may still be degraded by agricultural activity in the surrounding landscape.

Even with conservation policies to protect wetlands, we continue to witness wetland loss and degradation (Clare and Creed 2014; Davidson 2014). In the United States, for example, there are no federal policies that manage farming practices around wetlands (Johnston 2014; Mitsch and Gosselink 2015), though farmers require permits for activities that occur within a wetland under Section 404 of the Clean Water Act (U.S. EPA 2017). Similarly, the Albertan wetland policy (Government of Alberta 2013) offers legal protection to wetlands and introduces innovation in shifting the focus of management from an area-basis to a function-basis. The Albertan wetland policy fails to, however, provide protective buffers around wetlands (Government of Alberta 2013). Consequently, despite legal protections and regulation of activities that occur within wetland boundaries, wetland integrity and function may be compromised by adjacent human activities through connections linking wetlands to their catchments and beyond (Jones et al 2018; Kraft et al 2019).

Compromised wetland integrity may endanger bird populations because they are sensitive to both changes in wetland condition and landscape structure. For example, Mensing et al. (1998) found that, out of six taxa surveyed, birds were the best indicator of landscape condition surrounding small-stream riparian wetlands. Bird diversity and richness were highly correlated with the extent of cultivated land, wetland and forest cover within 500 and 1000 metre (m) radii (Mensing et al. 1998). These findings are echoed in research in Alberta’s Parkland region, which concluded that bird community integrity in shallow open-water wetlands was sensitive to road density, forest cover, and the amount of other wetland habitat within 500 m (Rooney et al. 2012).

Most research on the drivers of bird composition in wetlands have focused on permanently-ponded wetlands. Yet, temporarily-to semi-permanently-ponded wetlands also comprise high-value bird habitat, especially for breeding and brood-rearing birds (Burger 1985; Hands et al. 1991). Small, isolated wetlands sustain metapopulations (Semlitsch and Bodie 1998), and they are invaluable habitat for terrestrial, facultative, and obligate birds because 1) there are lower occurrences of mammalian predators (Burger 1985); 2) there are interspersions of mudflats that allow birds to dabble (Osborn et al. 2017), which allows them to feed while remaining alert for predators (Pöysä 1986); 3) macroinvertebrate prey are abundant and diverse (Zimmer et al. 2000; Gleason and Rooney 2017); and 4) the absence of fish improves food availability for birds (Zimmer et al. 2001). For example, Shealer and Alexander (2013) reported that insectivorous Black Terns (*Chlidonias niger*), which nest in more-permanently flooded wetlands, commonly forage in temporarily-flooded wetlands up to 4 kilometres (km) away. Since wetlands that dry up during the breeding season provide additional foraging opportunities for birds and refuge from predators, they are valuable bird habitat.

Since land-use is known to influence bird habitat selection (Ballard et al. 2014), we anticipate that bird community composition and guild structure in prairie pothole wetlands will be sensitive to the extent of agricultural activity in adjacent lands. We seek to determine which local- and landscape-level habitat variables are most predictive of bird assemblage occurrence in prairie potholes of Alberta. If birds are sensitive to agricultural activity in the surrounding landscape, it raises concerns that existing wetland policy that fails to provide buffer protections surrounding wetlands, may fail to protect bird communities and the important ecological services they provide. Furthermore, we evaluate the dependency of these predictions on wetland-dependent birds, including shorebirds, wetland-dependent songbirds and waterfowl. We asked 1) if there are distinct assemblages of birds occupying these wetlands, 2) if so, what habitat traits at the local- and landscape-level are predictive of assemblage occurrence; and 3) whether bird assemblages could be used to indicate the level of agricultural disturbance affecting a prairie pothole wetland.

## Methods

### Study Area

Our study region encompasses the Parkland and Grassland natural regions of Alberta (Fig. 1). In this semi-arid climate, evapotranspiration rates exceed annual precipitation (Downing and Pettapiece 2006; Millett et al. 2009), but depressions created by glaciation nonetheless give rise to a high density of small wetlands known as prairie potholes. The Parkland is cooler and moister, supporting a mosaic of deciduous forest and grassland. The Grassland is warmer and too dry for most trees (Downing and Pettapiece 2006).

**Fig. 1.**
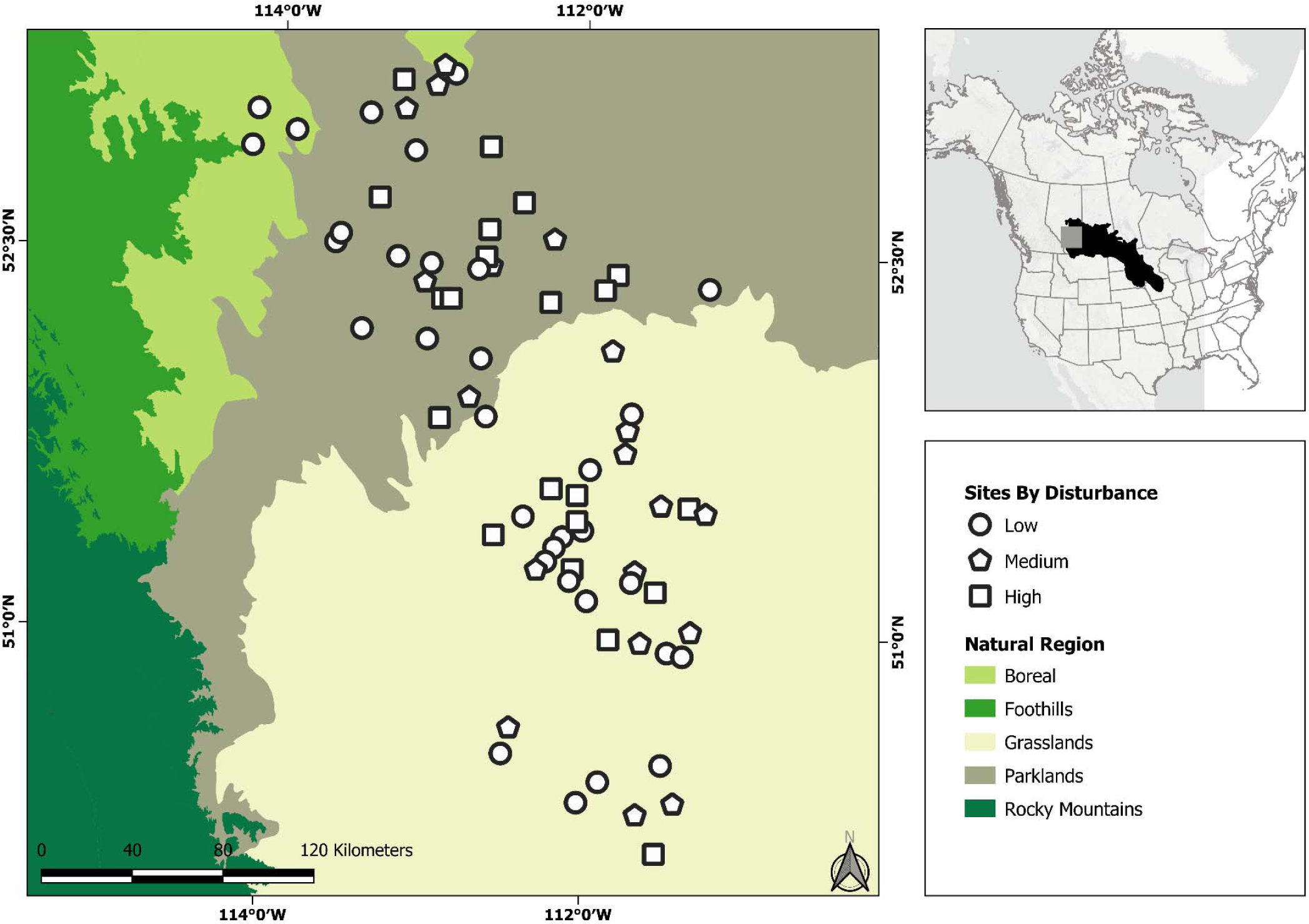
A map of study our region in the northern prairie pothole region. Our 72 wetland sites occupied both the Grasslands and Parklands region, belonging to temporary (*n*=11), seasonal (*n*=18), semi-permanent (*n*=10), and permanence (*n*=9) permanence classes.

### Study Design

We selected 72 natural wetlands than ranged in permanence class from temporarily-ponded to permanently-ponded (*sensu* Stewart and Kantrud 1971) and were evenly distributed among six, randomly selected sub-watersheds (three in each Natural Region – Grasslands and Parklands) of glaciolacustrine or glaciofluvial-derived surficial geology (Fig. 1). Our selected wetlands reflected the frequency distribution of wetland sizes within each sub-watershed, based on the Alberta Merged Wetland Inventory (Government of Alberta 2014a), and so were generally small (mean size 0.66 ± SE 0.07 ha). To guard against spatial dependencies, sites were spaced a minimum of 3.5 km apart. Independent of their hydroperiod, wetlands were selected to span a gradient in the extent of agricultural activity in the surrounding landscape (i.e., the percentage cropping, haying, and pastureland covers within 500 m buffers around each wetland’s perimeter). Land cover data were derived from the Agriculture and Agri-Food Canada Annual Crop Inventory Data (AAFC 2015) and supplemented with information from the provincial Grassland Vegetation Inventory (Government of Alberta 2014b). We used a 500 m buffer because this scale has been reported as the most influential of bird community integrity in permanently-ponded wetlands in Alberta’s Parkland (Rooney et al. 2012). Due to the level of agricultural activity in the Grassland natural region (Downing and Pettapiece 2006), where over 70% of land is privately owned (AEP 2011), truly pristine reference sites are scarce. Consequently, we classified wetlands with less than 25% agricultural land cover as being in the least disturbed condition (*sensu* Stoddard et al. 2006) and used these as a reference condition against which medium (25-75% agricultural cover) and high disturbance (greater than 75% agricultural cover) wetlands could be compared.

### Bird Surveys

Bird surveys were conducted by pairs of observers in 2014 and 2015, following the method described in Wilson and Bayley (2012). In brief, surveys comprised a 10-min visual survey followed by an 8-minute acoustic fixed-radius point count survey, with a radius of 100 m. Because most wetlands were less than 1 ha, a single 100-m radius point count covered the entire wetland. Larger sites were surveyed from two-point count locations, providing they could be positioned at a minimum of 200 m apart, in which case counts were summed to reflect the wetland as the sample unit.

Surveys were conducted twice at each wetland during the breeding season (May 19^th^ – June 24^th^) to account for any temporal partitioning of breeding activity among species within the general breeding season. Consequently, we summed counts across surveys. Generally, birds in our study region sing and call between sunrise and 11:00 am (Farr et al. 2012). Thus, all surveys were restricted to this time period.

All birds visually observed foraging or nesting or heard singing or calling at the site were enumerated and identified to species. Bird identifications followed the American Ornithologists Union Standard Information used to determine guild membership of bird species, such as feeding traits, preferred habitat, and migration patterns, were retrieved from Birds of North America Online (CLO 2014). We distinguish between this complete bird assemblage and the subset of birds observed using the marsh that are classified as wetland-dependent species (Online Resource 1); only these wetland-dependent species were included in our subsequent analyses.

### Local-level Habitat Characterization

We surveyed the vegetation at each wetland during peak aboveground biomass between late July and August. First, we used a sub-meter accuracy GPS (Juno Trimble T41; SXBlue II GPS/GNSS Receiver) to delineate the wetland boundary such that the perimeter of the wetland lay where vegetation transitioned to <50% cover by wetland-obligate plant species. Next, we sub-divided the wetland into zones based on vegetation form (woody vegetation, drawdown, ground cover, narrow-leaved emergent, broad-leaved emergent, robust emergent, open-water area) and the associated dominant or co-dominant macrophyte species. These vegetation zones were delineated in the same manner as the wetland and their area calculated in the field to inform quadrat-based sampling intensity. Each vegetation zone was then characterized by a minimum of five 1 m^2^ quadrats. If a zone was larger than 5000 m^2^, we added one quadrat per 1000 m^2^. Finally, we estimated the mean percent cover among plots and then relativized our estimates to 100%, for a site-level estimate of vegetation cover.

In addition to vegetation surveys, we monitored abiotic variables known to influence bird habitat selection. From May and September 2014, we measured water depth using staff gauges, providing ponded water remained in the wetland. This was used to calculate the wetland’s maximum water, minimum water depth, and seasonal amplitude (maximum depth minus minimum depth).

### Statistical Analysis

Our analysis objectives were 1) to test whether the birds grouped into distinct assemblages; and 2) to determine which local- and landscape-level variables were most predictive of bird assemblage occurrence by developing a model to classify wetlands in terms of their expected bird assemblage based on local- and landscape-variables, with particular emphasis on the level of agricultural activity surrounding each wetland.

We used a square-root transformation and relativized our wetland-dependent bird count data by the maximum value in each column to improve multivariate normality and reduce the influence of numerically-dominant species. To reduce data sparsity, we removed rare avian species (<2 occurrences out of 72 wetlands). Following the recommendations of McCune and Grace (2002), we used a Bray-Curtis dissimilarity measure to characterize distances in species space of community composition among our wetlands.

### Cluster & Indicator Species Analysis

We used a cluster analysis to identify distinct wetland-dependent bird assemblages among our sites. We used a hierarchical agglomerative polythetic process for the cluster analysis using the cluster package (Maechler et al. 2018) in R statistical software (R Core Team 2017). For the cluster analysis, we specified a flexible beta linkage method (beta = −0.250) and used the Bray-Curtis distance measure, based on the recommendations of McCune and Grace. (2002). Also using the cluster package, we then pruned the dendrogram iteratively, varying the number of groups among sites from two to 20.

We used an indicator species analysis (ISA) to determine the optimal number of wetland-dependent bird assemblages among our sites as the number of groups generating the smallest average *p*-value across indicator species. As described in Dufrêne and Legendre (1997), this analysis estimates the indicator value of each species based on their relative abundance and frequency in each group and assigns a measure of statistical significance using a Monte Carlos method with 4999 permutations. For the ISA, we used the labdsv package in R (Roberts 2016), and the site-group memberships (overall number of assemblages among our sites) derived from the trimmed dendrogram. In ISAs, because groups with one sample unit must be excluded from analysis (Peck 2010), we limited our analysis to site-group memberships with at least two sites per group.

### Visualizing Community Composition

To visualize how wetland-dependent bird communities are related to the local and landscape variables, we ran nonmetric multidimensional scaling ordinations (NMDSs) on our two bird matrices. We used the NMDSs to visualize how 1) the local and landscape variables in the final classification and regression tree (CART) model (described below) were related to community composition, and 2) functional traits were related to each wetland-dependent bird assemblage identified in the ISA. We used the vegan package to implement the NMDSs (Oksanen et al. 2017) in R statistical software (R Core Team 2017).

After implementing each NMDS, we used vector overlays to visualize how species counts (r^2^ > 0.2 with at least one axis) and counts of species possessing various functional traits aligned (r^2^ > 0.1 with at least one axis) with major trends in bird community composition. We symbolized sites by assemblages identified in the combined cluster analysis and ISA.

### Classification and Regression Tree

Finally, we developed a classification and regression tree to predict which wetland-dependent bird assemblage would occur at a marsh, using a combination of local- and landscape-level data. In our case, the classification and regression tree partitions the wetlands based on local- and landscape-level characteristics to create nodes of wetlands such that the deviance between node membership and bird assemblage cluster is minimized. We used local-level (size, percentage cover of woody, robust emergent and broad leave plants, maximum water depth) and landscape-level variables (percentage cover of grassland, forest and shrubs, water and wetlands, cropland and human-related land use within a 500 m radius) that would be critical in influencing the functional traits of wetland-dependent birds present in a wetland, as predictors in the classification tree.

We used the “tree” package (Ripley 2016) in R statistical software (R Core Team 2017), to implement the classification and regression tree. The classification tree implements binary recursive partitioning, using the deviance index described in Breiman et al. (1984) to estimate impurity for splitting, and stops splitting when the terminal node passes a size threshold for the number of wetlands included (Ripley 2016). Next, we used k-fold cross-validation to prune the tree, where k = 10, which was based on cost-complexity as measured by deviance. We also used the “tree” package to determine the number of misclassifications for the overall tree, as well as the number of misclassifications at each node. Because our small sample size could contribute to unstable k-fold cross-validation errors with increasing tree size, we repeated the test 100 times and found the mean and standard error across iterations.

We used goodness of fit tests to measure if our classification and regression tree predictions differed from the groups generated by the combined cluster analysis and ISA. Using the DescTools package (Signorell 2017) in R, and a Williams correction for our small sample size, we performed a G-Test. Next, we used the caret package (Kuhn 2017) in R to examine whether there was strong agreement between our classification and regression tree predictions and ISA assemblages, using kappa statistics.

## Results

### Cluster & Indicator Species Analysis

We differentiated five distinct wetland-dependent bird assemblages (dendrogram in Fig. 2; indicator values listed in Table 1), using agglomerative hierarchical clustering and ISA. We assigned each assemblage a name reflecting the life history traits of the birds that were the strongest indicators of the assemblage (indicator values listed in Table 1). Only a few species were considered significant indicators of the five wetland-dependent bird assemblages. Note that all but one wetland-dependent bird assemblage had at least one significant indicator species that was both faithful and relatively exclusive to that assemblage of birds. The exception is the Hummock Nesters (Table 1), which was the first assemblage to merge with another (Pond and Reed Associates) at a Bray-Curtis dissimilarity value of about 0.8 during cluster analysis. The strongest indicator species for the Hummock Nesters was Wilson’s Snipe (*Galliango delicata*), with an indicator value of 25.35 (p = 0.100). A list of indicator values of all bird species included in the cluster and indicator species analyses is presented in Table 1.

**Table 1.**
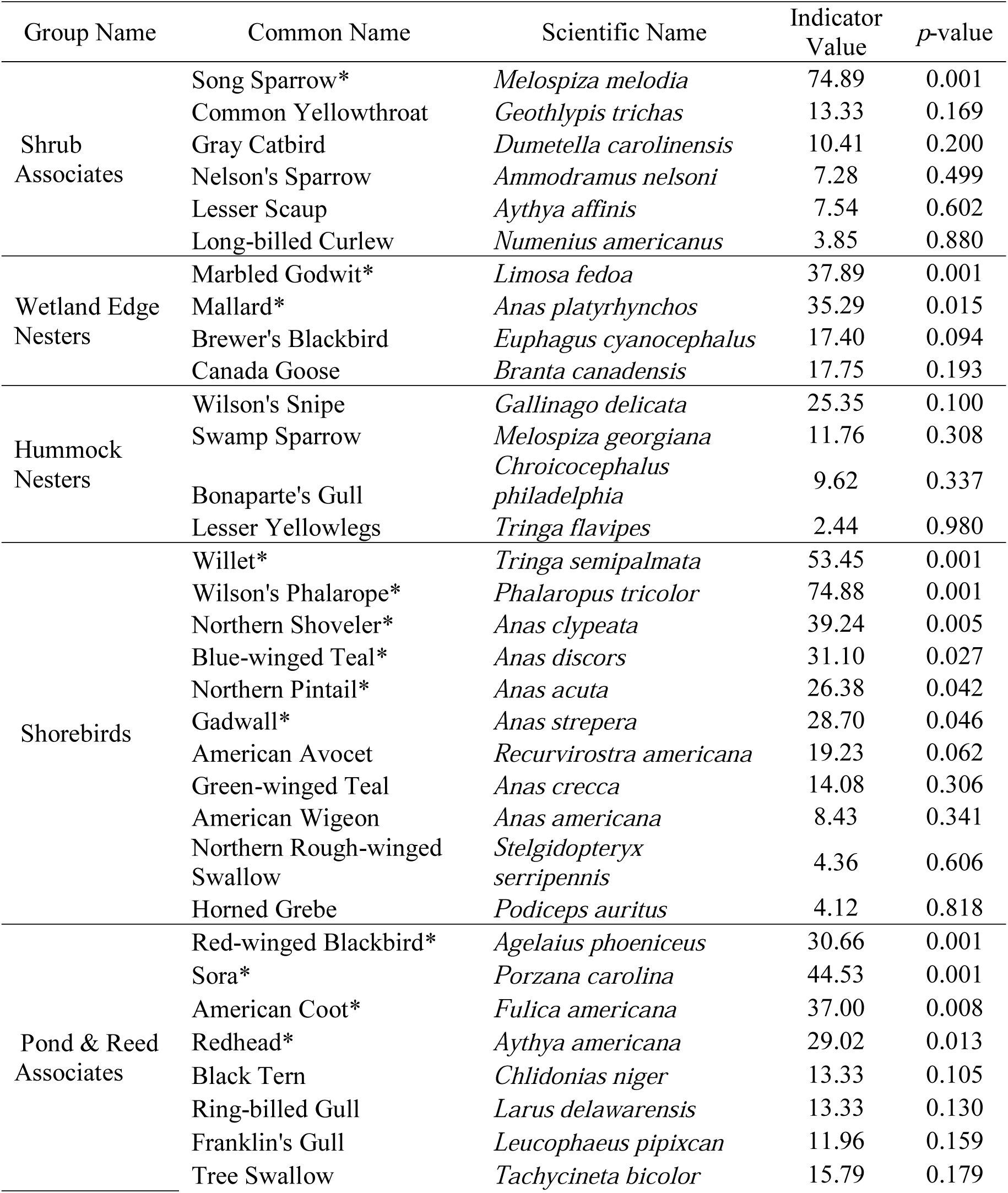
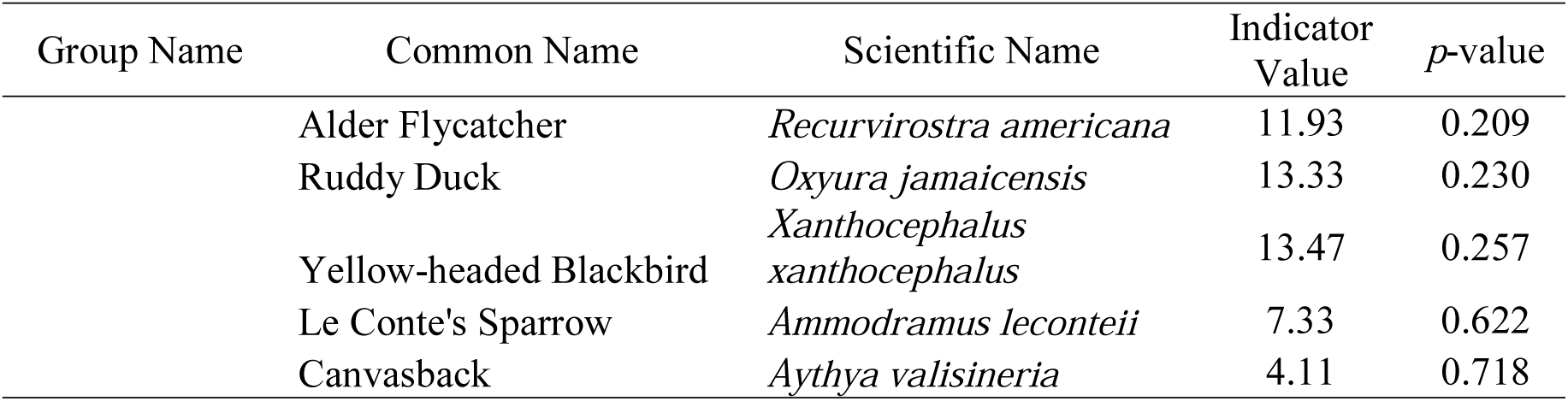
This table provides indicator values for species belonging to each of the five bird assemblages identified via cluster analysis of the dataset including only birds categorized as wetland-dependent species. Each species is grouped under the assemblage for which it had the highest indicator value, and the table includes all 38 species, regardless of whether it was a significant indicator of an assemblage. However, only 13 species were significant indicators (p<0.05), indicated by “*”. The associated *p*-value indicates the probability that an indicator value that large could be obtained from the data by chance alone. Note that the Hummock Nester assemblage was the first assemblage to merge (into the Pond and Reed Associates assemblage) during agglomerative clustering analysis (Figure 2) and had no significant indicators.

**Fig. 2.**
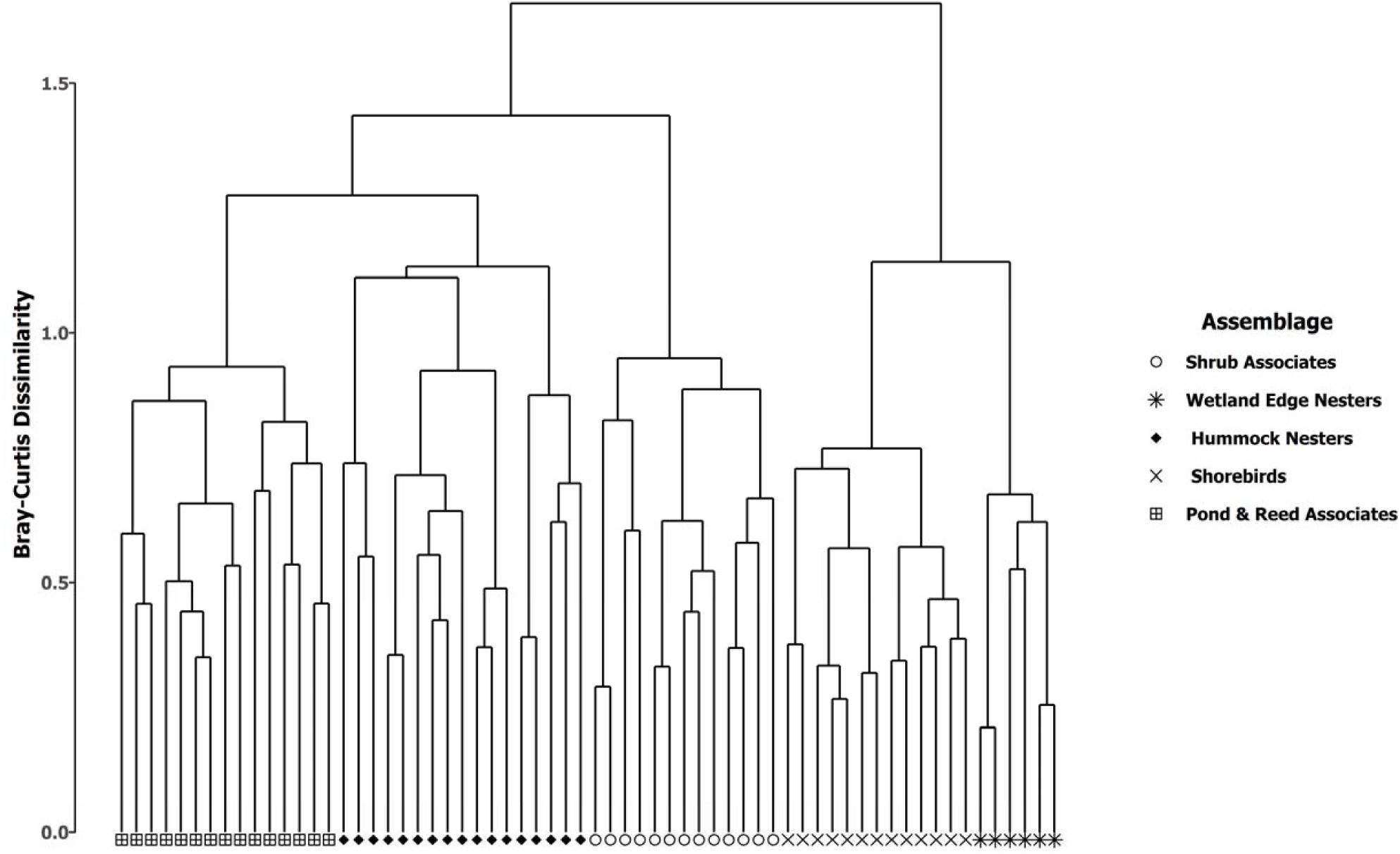
Dendrogram from agglomerative hierarchical clustering, where group membership was derived from indicator species analysis for birds categorized as wetland-dependent species only. Birds categorized as terrestrial species were excluded from this analysis (see Online Resource 1). Symbology of the sites at the tips of the dendrogram reflects the optimal dendrogram pruning level, determined using indicator species analysis. Group names were based on the life history traits of the dominant indicator taxa for each group.

### Visualizing Community Composition

Based on an assessment of the marginal decline in stress with increasing dimensionality, we concluded that a three-dimensional solution was optimal for both our NMDS ordinations. The NMDS, final stress was 18.58, and the NMDS converged in fewer than 20 runs.

The abundance of wetland-dependent birds differed among assemblages, based on their nesting or habitat preferences (Fig. 3). Unsurprisingly, the Shrub Associates assemblage supported shrub species [e.g. Gray Catbird (*Dumetella carolinensis*)]. However, both shorebird [e.g. Willet] and non-shorebird species [e.g. Mallards (*Anas platyrhynchos*)] were associated with the Wetland Edge Nester assemblage (Fig. 3C; 3D). Wetlands classified as supporting the Shorebird Assemblage contained abundant shorebird and ground nesting species (Fig. 3A; 3B). The Hummock Nester-classified wetlands shared species with all assemblages except the shrub associates (Fig. 3A; 3B), and they were not strongly associated with any bird nesting or habitat preferences (Fig. 3C; 3D). Conversely, only marsh (e.g. Sora) or pond species (e.g. American Coot) were associated with the Pond and Reed Nesters assemblage (Fig. 3A; 3C).

**Fig. 3.**
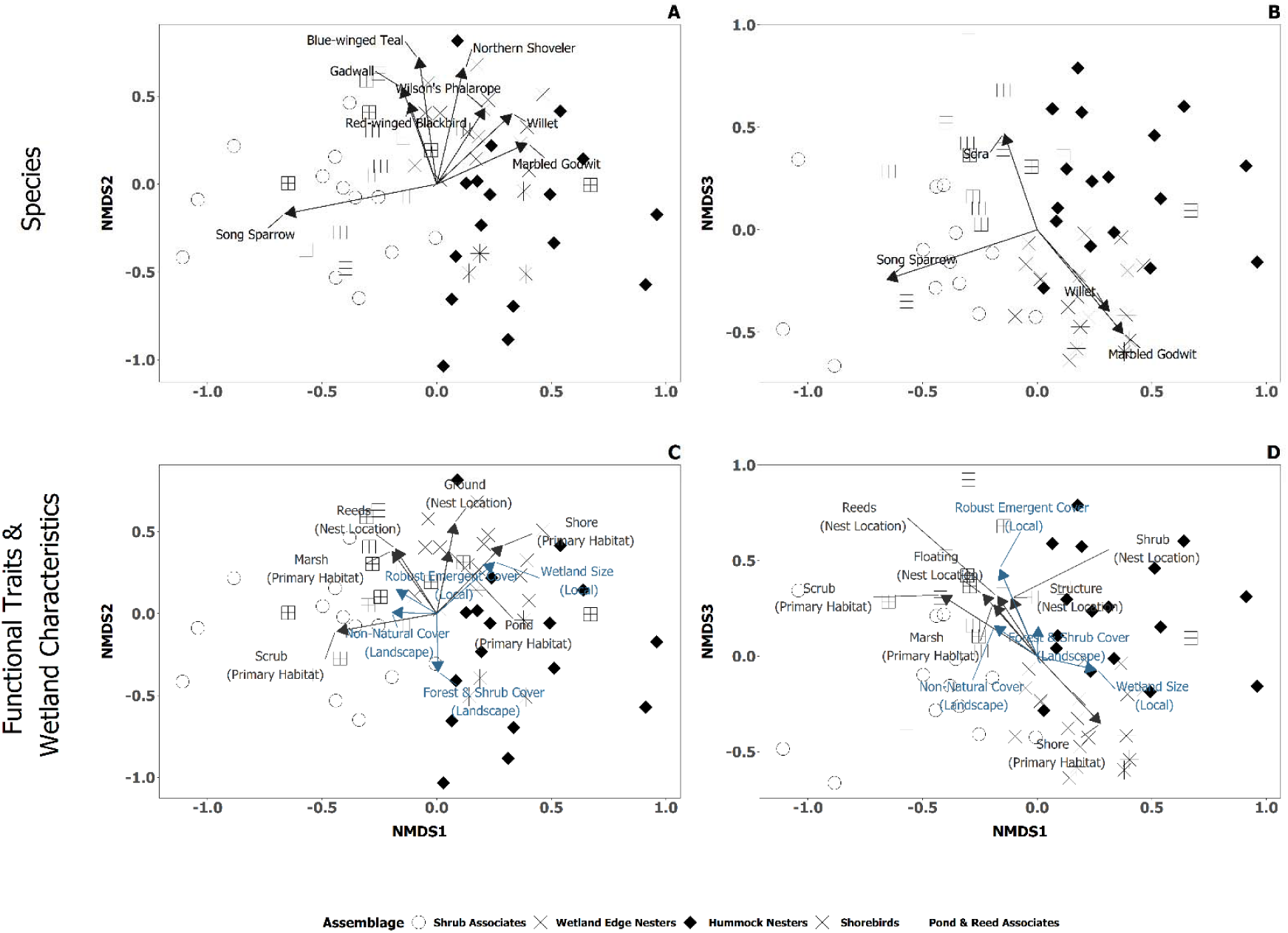
Plot of the nonmetric multidimensional scaling ordination for wetland-dependent birds, for both axis one and two (A & C) and two and three (B & D). Site assemblages are the result of our paired agglomerative hierarchical clustering and indicator species analyses. We estimated vectors from correlations between both axis and the 1) abundance of wetland birds (A & B), 2) abundance of wetland birds by functional traits (A & B), 3) percentage of non-natural cover and forest and shrub cover in the landscape (C&D) and 4) wetland size and percentage cover of robust emergent vegetation in a given wetland (C & D). Species vectors shown had r^2^ values greater than 0.2 for both axes; for functional traits, r^2^ values were greater than 0.1. Functional trait vectors indicate a birds’ nesting location and primary habitat (black), while wetland characteristics vectors indicate both local and landscape wetland characteristics (blue).

The NMDS axes were indicative of various local- and landscape-scale wetland characteristics. Axis one in the NMDS reflected a disturbance gradient (Fig. 3C; 3D), wetlands differed in which Natural Region they were located along axis two (Fig. 3C) and axis three reflected a hydroperiod gradient (Fig. 3D).

### Classification and Regression Tree

#### All Birds

Using a combination of local and landscape-level variables (comprehensive list of variables in Online Resource 2), we predicted which of the bird assemblages would occur at a given wetland. The classification tree had ten terminal nodes (Fig. 4), with low total residual deviance (12.48) and a misclassification error rate of 27%. Based on 10-fold cross validation error, we trimmed the tree from ten (cross-validation error = 60%) to eight (cross-validation error = 59%) terminal nodes (Table in Online Resource 3). The pruned tree had a marginally higher total residual deviance (12.94), but the same misclassification error rate (27%).

**Fig. 4.**
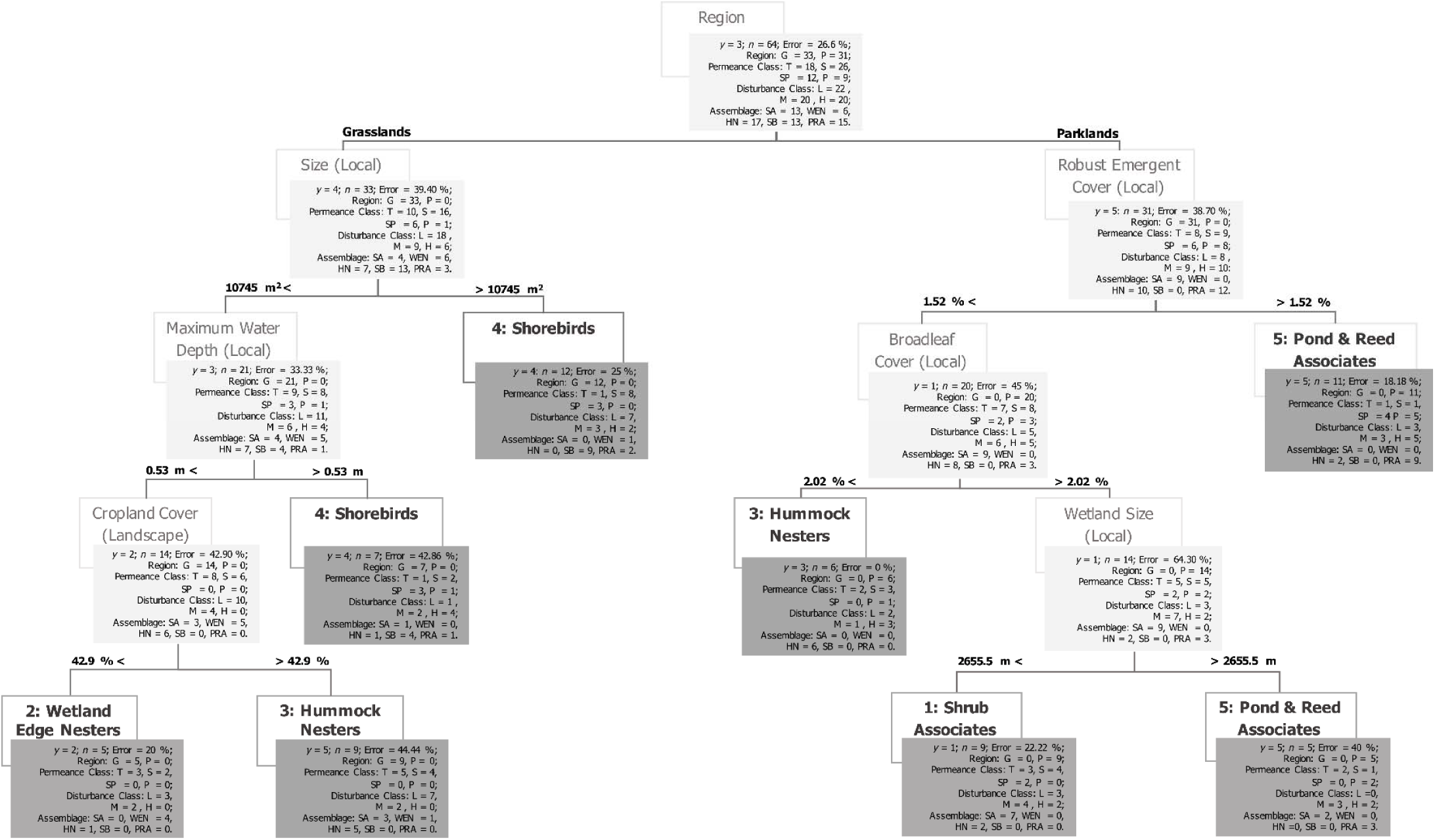
Classification and regression tree using a combination of local, landscape-level, and regional variables to predict a bird’s assemblage type. Assemblages are derived from cluster analysis carried out on the abundance of wetland birds we observed using the wetlands. Terminal nodes indicate the assemblage predicted to occur at this subset of wetlands by our classification tree. For each node, we present the 1) predicted assemblage; 2) misclassification error rate; and 3) the number of sites per region (G – Grassland and P – Parkland), permanence class (T – Temporary, S – Seasonal, SP – Semi-Permanent and P – Permanent), disturbance class (L – Low: >25 %, M – Medium: <25<75 % and H – High: <75 % non-natural cover), and observed assemblage (SA: Shrub Associates, WEN: Wetland Edge Nesters, HN: Hummock Nesters, SB: Shorebirds and PRA: Pond & Reed Associates).

The pruned model predicted all five assemblages. The model predictions had strong agreement with the assemblages from the ISA (kappa = 66%). More, any differences between the classification tree predictions and the observed assemblages were not statistically significant (G-Test: G = 10.19, df = 63, *p*-value =1.00).

The Wetland Edge Nesters and Shorebirds assemblages were the most distinct (in local and landscape characteristics), occurring in a single terminal node (Fig. 4). The other assemblages each occurred in two terminal nodes but differed in the distances between nodes (Fig. 4). The Shorebirds assemblage was the third-most distinct assemblage, though predicted at different tree heights. The Pond and Reed Nesters had the furthest vertical distance between the nodes. However, the Hummock Nester-classified wetlands were predicted in both regions, suggesting they were the least distinct assemblage.

Misclassification error rates were moderate (0 – 46%) (Table 2). Error rates were highest for adjacent assemblages (e.g. Wetland Edge Nesters vs. Hummock Nesters), supporting birds with similar foraging and nesting preferences. Conversely, error rates were low for assemblages that were restricted to a region (e.g. Shorebirds vs Pond and Reed Nesters) (Fig. 4; Table 2).

**Table 2.** Contingency table comparing the observed classification of wetland sites based on their comprehensive wetland-dependent bird assemblage and the predicted assemblage membership for each site. Predictions were based on a classification tree based on a set of local, landscape-level, and regional habitat characteristics (full list in Online Resource 2).

|  |  | Observed |  |  |  |  |
| --- | --- | --- | --- | --- | --- | --- |
|  |  | Shrub Associates | Wetland Edge Nesters | Hummock Nesters | Shorebirds | Pond & Reed Associates |
| Prediction | Shrub Associates | 7 | 0 | 2 | 0 | 0 |
|  | Wetland Edge Nesters | 0 | 4 | 1 | 0 | 0 |
|  | Hummock Nesters | 3 | 1 | 11 | 0 | 0 |
|  | Shorebirds | 1 | 1 | 1 | 13 | 3 |
|  | Pond & Reed Associates | 2 | 0 | 2 | 0 | 12 |

Local wetland characteristics were most predictive of assemblages (Fig. 4). Similar to our analysis on all birds using a wetland, Natural Region was the strongest predictor of assemblages. Apart from Hummock Nesters and Wetland Edge Nesters, the assemblages were predicted by proxies for wetland hydroperiod (e.g., size, depth) and vegetation characteristics (e.g., robust emergent vegetation cover, broadleaf vegetation cover).

## Discussion

Although wetland policy across North America aims to protect wetlands, neither American nor Canadian policy limits what activities can take place in the immediate landscape surrounding wetlands. If wetland bird communities are sensitive to adjacent land cover/land use, then existing policy may be incapable of conserving wetland functions without incorporating some buffer protections. Conversely, if avifauna are sensitive only to in situ wetland conditions such as hydrology and vegetation structure, then buffer protections should not be necessary to conserve bird community function and integrity. Based on prior research (e.g., Rooney et al 2012; Anderson and Rooney 2019), we predicted that bird community composition and guild structure in wetlands of the Grassland and Parkland Natural Regions of Alberta would be sensitive to the proportion of agricultural land cover in the surrounding landscape. We found some support for this prediction – one of our five wetland-dependent bird assemblages were absent from wetlands in agriculturally-dominated landscapes. We attribute this to differences in the nesting behaviors of this assemblage – they nested in upland habitat and consequently selected for landscapes with more natural land cover, mainly grassland. Conversely, dabblers, divers, and waders were indicative of assemblages in landscapes with more agricultural activity and longer hydroperiods. Consequently, waterfowl and shorebirds are less sensitive to this land conversion, and so come to dominate wetlands situated in agricultural landscapes. However, the importance of surrounding land cover in predicting which wetland-dependent assemblage would occur at a wetland was less evident than we anticipated.

The most significant predictor of bird assemblage occurrence was the Natural Region that the wetland fell in – Parkland vs. Grassland. The Grassland and Parkland natural regions differ in both their landscape- and local-level characteristics. At the landscape-level, the Parkland supports copses of aspen forest and more shrubland than the Grassland (Downing and Pettapiece 2006). Further, while there is more cropland in the Parkland, pastureland and haying are more common in the Grassland (Downing and Pettapiece 2006). Also, because of differences in climate, we observe a higher abundance of wetlands with longer hydroperiods in the Parkland (Government of Alberta 2014a). In the more arid Grassland, the magnitude of difference between potential evapotranspiration and precipitation is larger, resulting in the dominance of shorter-hydroperiod wetlands (e.g., temporary and seasonal). Thus, we likely find Natural Region to be a strong predictor of avian assemblage occurrence because of the preference of some bird species for shorter-hydroperiod wetlands in mixed-grass prairie typical of the Grassland (e.g., Shorebirds) versus the preference of other bird species for longer-hydroperiod wetlands in landscapes with more shrubland and forest typical of the Parkland (e.g., Pond/Reed Associates).

The composition of the landscape surrounding a wetland is not strongly predictive of which wetland-dependent assemblage we find occupying a wetland. Local-level factors such as hydroperiod (wetland permanence class, depth and size) and vegetation characteristics (percentage cover of broadleaf and robust emergent plants) were important predictors of assemblage occurrence. An explanation for this influence of local-level factors is that both wetland hydroperiod and vegetation dictate food availability (Lantz et al. 2011) and nesting opportunities for wetland-dependent birds (Kantrud and Stewart 1984). Because many wetland-dependent birds have feeding behaviors (e.g., diving, dabbling, and wading) tied to the availability of open water habitat, these local-level factors were critical in determining whether their habitat needs could be met. For instance, the Pond and Reed Associate and the Shorebird assemblages were distinguished using both the all birds and wetland-dependent bird datasets. These assemblages were characterized by birds that dive, dabble, and wade to feed. More, these assemblages were predicted to occur in wetlands that were deeper, larger, and had longer hydroperiods and nearly all their indicator species were ground, pond or reed nesters that nest in the wetland proper. Thus, we conclude that these assemblages were most sensitive to in situ factors about the wetland, rather than the character of its surrounding landscape.

Waterfowl and shorebirds may dominate wetlands in landscapes heavily influenced by agriculture not necessarily because they profit from cropping and grazing activities, but because species reliant on upland habitat for nesting are excluded. Both the Wetland Edge Nesters and Hummock Nesters assemblages were predicted to occur in deeper (>0.53 m), larger (>10745 m^2^) wetlands in the Grassland, but it was the Hummock Nester assemblage that occurred in wetlands with higher cropland activity in the surrounding landscape (>42 %). Consequently, species belonging to the Hummock Nester assemblage come to dominate these wetlands because their nesting habitat is still available when upland habitat is lost to agriculture, while wetland birds that typically nest in upland habitat are now unable to do so (e.g., species the Wetland Edge Nester assemblage). Similarly, Anderson and Rooney (2019) reported that significant differences in bird community composition between natural and restored wetlands in the Parkland region of Alberta were only evident when all birds were considered. They also reported that any difference in the composition of wetland-dependent birds were negligible because restored wetlands were similar to natural wetlands in their size, hydroperiod, and vegetation zonation, but differed significantly in terms of landscape context. Therefore, by using a more comprehensive bird survey data, we can develop bird-based wetland monitoring and assessment tools that reflect the community-wide impacts of land cover change on bird assemblage occurrence.

Our CART and NMDS can be useful tools in designing wetlands for wetland-dependent birds. Though the species pool did differ between Natural Regions, landscape composition can be important when designing wetlands for birds. For example, if a practitioner aimed to provide habitat for a Shorebird assemblage in the Grassland Natural Region, the wetland should be deep (> 0.53 m) or large (>10745 m^2^) (i.e., CART results) and have lower human activity (i.e., NMDS results). However, if the said practitioner was targeting Wetland Edge Nesters, the wetland can be smaller (<10745 m^2^) but should have moderate to low cropping activity in the landscape (<42.9 %).

## Conclusion

We show that, generally, wetland-dependent assemblages show poor sensitivity to agricultural activity. While waterfowl and shorebirds were sensitive to in situ properties of the wetland, such as water depth and wetland size or vegetation zonation patterns, edge-nesting birds were excluded from wetlands with higher cropping activity. Waterfowl and shorebirds seem to dominate wetlands in landscapes with more agricultural activity because other avian species are excluded, despite being at greater risk of predation in these landscapes (Emery et al. 2005). SWhen designing wetlands for use by these wetland avifauna, our concurrent analyses using a CART and NMDS are useful tools in determining the landscape context and wetland characteristics suitable for assemblages that may be the target in restoration policy.

## Acknowledgements

Funding for this research was provided by Alberta Innovates grant #2094A. We are grateful to Dr. Derek Robinson who assisted with site selection and land cover analysis. We are also grateful to Drs. Stephen Murphy and Roland Hall who provided feedback on an early draft of this manuscript. We thank Daina Anderson, Brandon Baer, Matt Bolding, Graham Howell, Adam Kraft, Jennifer Gleason and Nicole Meyers for collecting the field data. Finally, we thank Dr. Erin Bayne for supplying the automated recording units, which were used to verify auditory surveys.

---

**Online Resource 1.** List of species included in our analysis. Species listed as wetland obligate – wet are known to nest in wetlands (i.e., shorebirds and waterfowl), while species listed as wetland obligate – dry (e.g., Bonaparte’s Gull, Brewer’s Blackbird) may nest in forests or grasslands at the wetland edge.

| Common Name | Scientific Name | Wetland-Dependent Species |  |
| --- | --- | --- | --- |
|  |  | Wetland Obligate – Wet | Wetland Obligate – Dry |
| Alder Flycatcher | *Recurvirostra americana* |  | * |
| American Avocet | *Recurvirostra americana* | * |  |
| American Coot | *Fulica americana* | * |  |
| American Wigeon | *Anas americana* | * |  |
| Baltimore Oriole | *Icterus galbula* |  | * |
| Barn Swallow | *Hirundo rustica* |  | * |
| Black-billed Magpie | *Pica hudsonia* |  | * |
| Black Tern | *Chlidonias niger* | * |  |
| Bonaparte's Gull | *Chroicocephalus philadelphia* |  | * |
| Brewer's Blackbird | *Euphagus cyanocephalus* |  | * |
| Blue-winged Teal | *Anas discors* | * |  |
| Canada Goose | *Branta canadensis* | * |  |
| Canvasback | *Aythya valisineria* | * |  |
| Clay-colored Sparrow | *Spizella pallida* |  | * |
| Common Yellowthroat | *Geothlypis trichas* |  | * |
| Eastern Kingbird | *Tyrannus tyrannus* |  | * |
| Franklin's Gull | *Leucophaeus pipixcan* | * |  |
| Gadwall | *Anas strepera* | * |  |
| Gray Catbird | *Dumetella carolinensis* |  | * |
| Green-winged Teal | *Anas crecca* | * |  |
| Horned Grebe | *Podiceps auritus* | * |  |
| Long-billed Curlew | *Numenius americanus* |  | * |
| Le Conte's Sparrow | *Ammodramus leconteii* |  | * |
| Lesser Scaup | *Aythya affinis* | * |  |
| Lesser Yellowlegs | *Tringa flavipes* | * |  |
| Marbled Godwit | *Limosa fedoa* |  | * |
|  |  | Wetland Obligate – Wet | Wetland Obligate – Dry |
| Mallard | *Anas platyrhynchos* | * |  |
| Nelson's Sparrow | *Ammodramus nelsoni* | * |  |
| Northern Pintail | *Anas acuta* | * |  |
| Northern Rough-winged Swallow | *Stelgidopteryx serripennis* |  | * |
| Northern Shoveler | *Anas clypeata* | * |  |
| Ring-billed Gull | *Larus delawarensis* | * |  |
| Redhead | *Aythya americana* | * |  |
| Red-eyed Vireo | *Vireo olivaceus* |  | * |
| Ruddy Duck | *Oxyura jamaicensis* | * |  |
| Red-winged Blackbird | *Agelaius phoeniceus* |  | * |
| Savannah Sparrow | *Passerculus sandwichensis* |  | * |
| Sora | *Porzana carolina* | * |  |
| Song Sparrow | *Melospiza melodia* |  | * |
| Swamp Sparrow | *Melospiza georgiana* | * |  |
| Tree Swallow | *Tachycineta bicolor* |  | * |
| Willet | *Tringa semipalmata* | * |  |
| Wilson's Phalarope | *Phalaropus tricolor* | * |  |
| Wilson's Snipe | *Gallinago delicata* | * |  |
| Yellow-headed Blackbird | *Xanthocephalus xanthocephalus* | * |  |

**Online Resource 2.** A summary of site, landscape-level and local characteristics of our study wetlands, based on the assemblages’ classification from the indicator species analysis for birds categorized as wetland-dependent species. Besides region, we only used local and landscape characteristics in the classification tree. When the variable was continuous, we present the mean value with standard errors. Otherwise, we present the number of sites belonging to the category. Variables with the “*” symbol are percentage cover estimates, either at the local-level, or within a 500-m buffer landscape surrounding the wetland.

| Assemblage |  | Shrub Associates | Wetland Edge<br>Nesters | Hummock<br>Nesters | Shorebirds | Pond & Reed<br>Associates |
| --- | --- | --- | --- | --- | --- | --- |
| **Site Characteristics** |  |  |  |  |  |  |
|  | *n* | 13 | 6 | 17 | 13 | 15 |
| **Permanence**<br>**Class** | Temporary | 4 | 4 | 6 | 2 | 2 |
|  | Seasonal | 6 | 2 | 7 | 8 | 2 |
|  | Semi-Permanent | 2 | 0 | 3 | 3 | 4 |
|  | Permanent | 1 | 0 | 1 | 0 | 7 |
| **Region** | Grassland | 4 | 6 | 7 | 13 | 3 |
|  | Parkland | 9 | 0 | 10 | 0 | 12 |
| **Disturbance**<br>**Class** | Low | 5 | 3 | 7 | 7 | 2 |
|  | Medium | 6 | 2 | 4 | 3 | 5 |
|  | High | 2 | 1 | 6 | 3 | 8 |
| **Local Characteristics** |  |  |  |  |  |  |
| **Maximum Water Depth (m)** |  | 0.44 ± 0.06 | 0.26 ± 0.05 | 0.42 ± 0.06 | 0.47 ± 0.05 | 0.62 ± 0.07 |
| **Wetland Size (m<sup>2</sup>)** |  | 3159.54 ± 961.17 | 4679.83 ± 2098.36 | 6030.29 ± 1907.40 | 14757.77 ± 2693.90 | 9292.07 ± 2340.74 |
| **\*Broadleaf Plants (%)** |  | 0.95 ± 0.39 | 5.22 ± 5.07 | 7.56 ± 3.15 | 3.28 ± 1.34 | 0.65 ± 0.25 |
| **\*Woody Plants (%)** |  | 0 ± 0 | 0 ± 0 | 0 ± 0 | 0 ± 0 | 0.04 ± 0.04 |
| **\*Robust Emergent Plants (%)** |  | 0.62 ± 0.57 | 0.22 ± 0.22 | 1.56 ± 1.00 | 0.79 ± 0.70 | 8.82 ± 2.76 |
| **Landscape Characteristics** |  |  |  |  |  |  |
| **\*Non-Natural (%)** |  | 38.64 ± 9.04 | 34.13 ± 15.46 | 41.13 ± 9.42 | 39.4 ± 11.38 | 66.23 ± 7.15 |
| **\*Cropland (%)** |  | 12.46 ± 5.63 | 53.26 ± 15.35 | 30.47 ± 9.49 | 20.09 ± 7.83 | 42.48 ± 9.31 |
| **\*Urban Exposed Lands (%)** |  | 1.90 ± 0.55 | 3.91 ± 1.25 | 1.69 ± 0.48 | 4.01 ± 1.79 | 3.22 ± 0.66 |
| **\*Pasture & Forage (%)** |  | 18.09 ± 5.44 | 3.67 ± 1.82 | 4.64 ± 1.42 | 10.17 ± 4.05 | 19.5 ± 5.57 |
| **\*Water & Wetland (%)** |  | 5.64 ± 1.73 | 5.32 ± 2.83 | 7.24 ± 1.96 | 5.07 ± 2.18 | 10.87 ± 2.29 |
| **\*Forest & Shrub (%)** |  | 19.89 ± 8.89 | 15.87 ± 8.93 | 22.34 ± 7.21 | 2.18 ± 1.85 | 6.89 ± 2.47 |

**Online Resource 3.** Cross-validation error for the classification tree for birds categorized as wetland-dependent species, based on the number of terminal nodes. We found the mean and standard error for cross-validation across 100 iterations.

| Terminal Nodes | Error |
| --- | --- |
| 10 | $59.74 \pm 0.49$ |
| 8 | $58.66 \pm 0.47$ |
| 7 | $60.63 \pm 0.51$ |
| 3 | $64.05 \pm 0.43$ |
| 2 | $76.25 \pm 0.44$ |
| 1 | $82.42 \pm 0.30$ |

## Notes

https://doi.org/10.5061/dryad.524cv34

### Competing Interest Statement

The authors have declared no competing interest.

